# Viruses.STRING: A virus–host protein–protein interaction database

**DOI:** 10.1101/396184

**Authors:** Helen Victoria Cook, Nadezhda Tsankova, Damian Szklarczyk, Christian von Mering, Lars Juhl Jensen

**Affiliations:** Novo Nordisk Foundation Center for Protein Research, University of Copenhagen; Center for non-coding RNA in Technology and Health, University of Copenhagen; Swiss Institute of Bioinformatics, University of Zurich

## Abstract

As viruses continue to pose risks to global health, having a better understanding of virus–host protein–protein interactions aids in the development of treatments and vaccines. Here, we introduce Viruses.STRING, a protein–protein interaction database specifically catering to virus-virus and virus-host interactions. This database combines evidence from experimental and text-mining channels to provide combined probabilities for interactions between viral and host proteins. The database contains 177,425 interactions between 239 viruses and 319 hosts. The database is publicly available at viruses.string-db.org, and the interaction data can also be accessed through the latest version of the Cytoscape STRING app.

## Background

Viruses are well known as global threats to human and animal welfare. Viral diseases such as hepatitis caused by Hepatitis C virus (HCV) and cervical cancer caused by Human papillomavirus (HPV) each cause more than a quarter of a million deaths worldwide each year [1]. Outbreaks also present an economic burden - the 2014 Ebola virus outbreak, cost 2.2 billion USD to contain [2], and the annual response to Influenza virus costs 5 times this amount in medical expenses in the US alone [3]. Climate change and changing land use patterns are causing humans and livestock to be exposed to novel viruses for which there are currently no vaccines or antiviral drugs [4]. This trend will continue as the habitats of vectors that carry arboviruses expand [5], and as humans continue to come into contact with wildlife, creating opportunities for zoonosis [6].

As obligate intracellular parasites, viruses act as metabolic engineers of the cells they infect as they commandeer the cell’s protein synthesis mechanisms to replicate [7]. Thus, it is important to study their interactions with host cells in order to understand their biology, especially how their disruption of the host protein–protein interaction network causes disease [8]. Antiviral drugs have been highly effective at preventing the progression of HIV infection to AIDS [9], however, the effectiveness of antiviral drugs can decrease over time due to the development of drug resistant viral strains [10, 11, 12, 13]. A more complete understanding of the host-virus protein-protein interaction network provides more potential viral drug targets, and also enables alternative strategies such as targetting host proteins to attenuate viral infection [14]. When available, vaccines are very effective at preventing diseases caused by viruses [15]; however, vaccines are not available for all viruses, including HIV-1 and HCV, and a universal Influenza vaccine is still elusive [16]. The development of modern vaccines such as subunit vaccines, which can be administered to immunocompromized patients, and which eliminate the chance that the vaccine could revert to an infectious virus [17], also hinges on understanding the protein–protein interactions between viruses and their hosts.

Novel protein–protein interaction (PPI) information is disseminated primarily in the scientific literature, but it is not always organized in ways that make it easy to find, access, or extract. Databases such as VirusMentha [18] and HPIDB [19] make strong efforts to organize virus–virus and virus–host PPIs into databases, where this information is available in an easily parsable format. However, with the volume of the biomedical literature growing exponentially at 4% per year [20], it is not feasible for human curators to thoroughly review all new publications to add any new evidence to curated databases [21]. Automated text-mining methods are thus required to get a comprehensive picture of what is already known about the viruses we study.

We have expanded the popular database STRING [22] to include intra-virus and virus-host PPIs. The STRING database has been in constant development for 15 years, and the current version includes protein interaction data for over 2000 species, however all the interactions are exclusively intra-species. In this work, for the first time, we include cross species interactions into the STRING database. The PPIs reported by STRING represent functional associations between proteins. These interactions are not limited to physical interactions, and may also include interactions such as transcription factor binding, or the interaction may represent the fact that the associated proteins appear in the same biological pathway. In this paper the terms “interaction” and “PPI” are used to refer to functional associations. STRING combines many different sources (channels) of information to give a confidence score that measures the probability that the interaction is true. In a similar fashion, we provide virus-related probabilistic interaction networks derived from text mining and experiments channels.

## Methods

### Text mining evidence

Text mining for virus species and proteins was conducted using the dictionary-based software described in [23], the same tool that is used for the STRING text mining pipeline. The dictionary for virus species was constructed from NCBI Taxonomy [24], with additional synonyms taken from Disease Ontology [25] and the ninth ICTV report on virus taxonomy [26] to give 173,767 names for 150,885 virus taxa. The virus protein dictionary was constructed from the 397 reference proteomes that were present in UniProt [27] on Aug 31, 2015. All virus protein names and aliases were expanded following a set of rules to generate variants. This gave 16,580 proteins with 112,013 names. This dictionary was evaluated against a benchmark corpus of 300 abstracts that were annotated by domain experts [28]. The host species and protein dictionaries were identical to those used during the text mining for STRING 10.5 [22]. The text mining was conducted over a corpus that contained the more than 26 million abstracts in PubMed [20], and more than 2.2 million full text articles. The interactions found by this method represent functional associations between the identified proteins.

### Experimental evidence

Experimental data for virus–virus and virus–host PPIs was imported from BioGrid [29], MintAct [30], DIP [31], HPIDB [19] and VirusMentha [18]. These virus–host interactions were scored and then benchmarked against a gold standard set derived from KEGG. This creates a mapping between the number of interacitons mentioned in a study and the probability that they are true interactions according to the benchmark set [32]. The interactions found by this method represent physical interactions.

### Transfer evidence

Orthology relationships were used to transfer interactions following the same protocol that STRING uses, which is briefly described here. Both virus and host orthology relations were taken from EggNOG 4.5 [33]. STRING transfers an interaction between two proteins of the same species to two orthologous proteins in another species as is shown in figure 1a, and exactly the same was done to also transfer virus–virus PPIs. For transfer of a host–virus PPI, three cases are possible and are illustrated in figure 1b-d. The known interaction between a virus protein and host protein could be transferred to an orthologous virus protein in a different virus species (panel b), to an orthologous host protein in a different host (panel c), or both cases simultaneously, to both a new virus and a new host (panel d). Transfer is made only between viruses and the hosts they are known to infect, ie we do not predict new host-virus pairs based on orthology.

**Figure 1:**
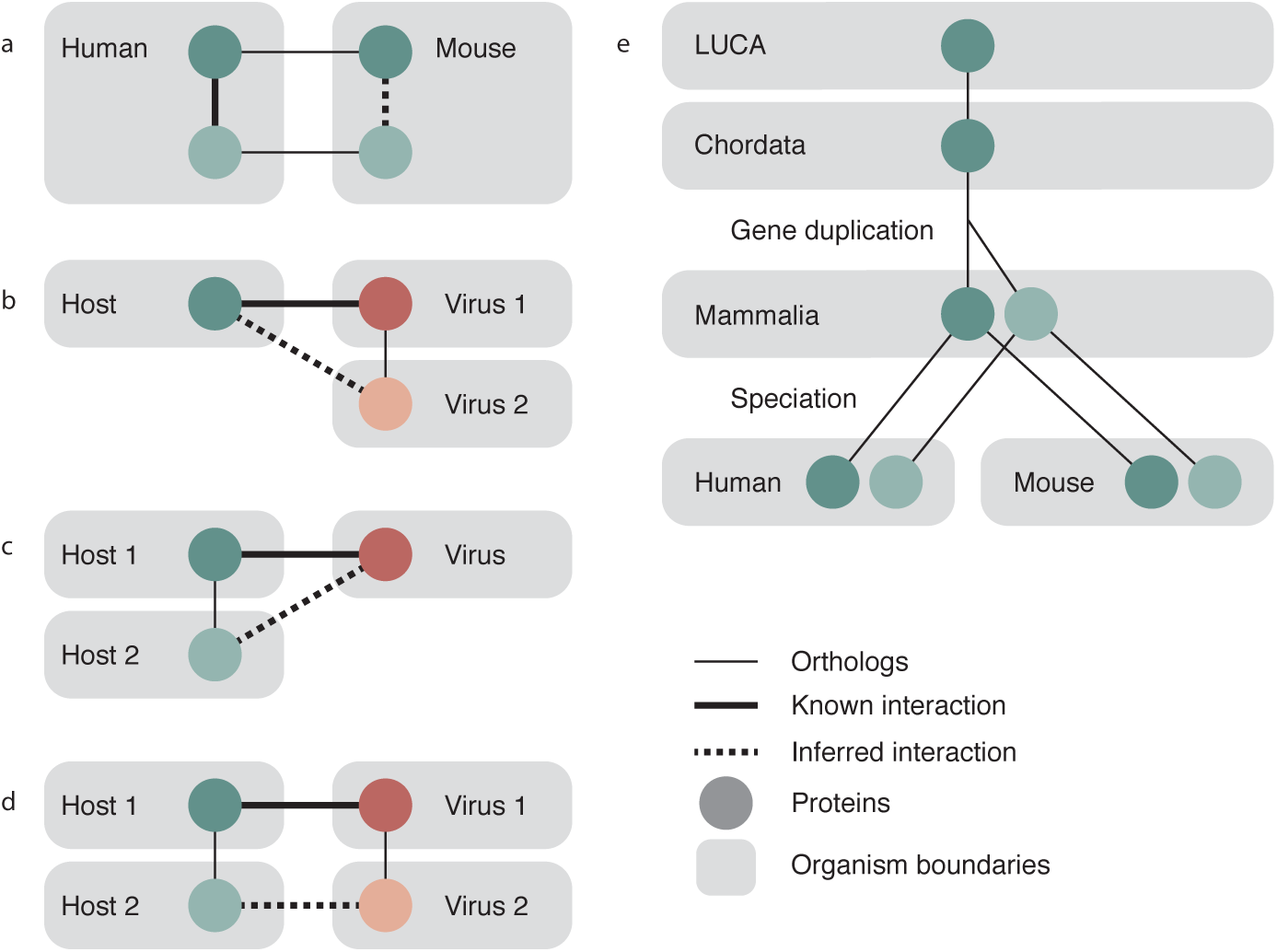
Orthology transfer in Viruses.STRING. STRING intra-species interactions are transferred between organisms as shown in panel a: an interaction between two proteins in species 1 (solid thick line) is transferred to two orthologous proteins in species 2 (dashed line). Orthology relationships are indicated by solid thin lines. This relationship is identical to transferring an interaction between two virus proteins of one virus species to two orthologous proteins in another virus species. Cross species interactions are handled as one of three cases — same host to closely related virus (panel b), same virus to closely related host (panel c), or both a new host and new virus (panel d). Panel e shows the evolutionary history of a gene that underwent a gene duplication event after the last common ancestor of chordata, but prior to the last common ancestor of mammals. There was subsequently a speciation event that resulted in the duplicated gene being present in both human and mouse. Orthology groups can be read by following the lines up the tree — at the level of mammalia, the light and dark genes are in separate orthology groups, but at higher levels, they are in the same orthology group.

The score assigned to the transfer of evidence is a scaled fraction of the score for the original interaction, proportional to how distant the recipient species is. Paralogs are considered to be orthologs for the purposes of calculating the score at level lower than the gene duplication, and the score is discounted if it is being used at a level higher than the gene duplication. For example, figure 1e shows three orthology levels (LUCA, Chordata, Mammalia) and illustrates a gene that has duplicated after Chordata but prior to the last common ancestor of all mammals. Further, there has been a speciation event after Mammalia, separating human and mouse into separate species. At the level of Mammalia, these two proteins are placed in different orthology groups, so any interactions that occur with the darker protein will not be transferred to interaction evidence for the lighter protein. However, at the level of Chordata, the light and dark proteins are in the same orthology group and so will both contribute their confidence to the resulting interaction. The contribution of these two proteins will be penalized since they are paralogs at a lower level. Although it is illustrated here for cellular organisms, this process is also applied to transfer involving viral orthology groups. The final transfer scores are then benchmarked the same way as the scores for the other channels.

## Results

We were able to identify 177,425 protein-protein interactions for 239 viruses. 77 of which are human viruses, and the remainder infect a total of 318 other hosts. The median number of proteins coded for by these viruses is 9. The majority of all types of interactions are between viruses and their hosts (as opposed to being intra-virus interactions), due to viral genomes encoding many fewer proteins than than their host genomes and thus having fewer potential interactions. In this and the subsequent analysis, interactions are counted per channel, disregarding their scores. Excluding orthology transfer, 89% of the interactions are derived from text mining evidence, and the remaining untransferred evidence comes from curated experimental databases. For 154 viruses, representing 19.8% of all evidence in the database, only text mining evidence is present. For 77 viruses, representing 77.4% of all evidence, all the experimental evidence is also supported by text mining evidence. The remaining 8 viruses, representing 2.8% of evidence, have more experimental evidence than text mining, and likely represent opportunities to improve the text mining dictionaries. Despite the large efforts of database curators, the vast wealth of information on PPIs is accessible only in the literature. Further, in addition to physical interactions, text mining will also uncover functional associations such as genetic interactions. As such, text mining provides a very important contribution to this database.

The top GO terms that are enriched in the set of 1835 human proteins that interact with any virus protein with a confidence of 0.5 or greater are shown in table 1. That this list includes terms such as viral process, protein binding and cell surface receptor signalling pathway provides a sanity check that the human protein partners in the found interactions are valid.

**Table 1:** Top GO terms by p-value that are enriched for the human proteins that interact with any virus protein.

| Number of genes | FDR p-value | GO term |
| --- | --- | --- |
| 549 | 2.28E-104 | positive regulation of macromolecule metabolic process |
| 718 | 5.96E-93 | positive regulation of cellular process |
| 452 | 3.02E-91 | cell surface receptor signaling pathway |
| 758 | 7.87E-91 | protein binding |
| 779 | 1.23E-90 | positive regulation of biological process |
| 539 | 1.61E-87 | positive regulation of cellular metabolic process |
| 459 | 3.86E-84 | multi-organism process |
| 277 | 4.56E-82 | innate immune response |
| 468 | 5.27E-82 | carbohydrate derivative binding |
| 595 | 1.99E-81 | response to stress |
| 240 | 2.07E-81 | multi-organism cellular process |
| 254 | 2.4E-81 | regulation of immune response |
| 239 | 2.55E-81 | viral process |
| 583 | 4.17E-81 | regulation of response to stimulus |

Orthology transfer gives a 2.7 times increase in the number of interactions with text mining results being more readily transferred than experimental results. A handful of well studied viruses (EBV, HIV-1, Influenza A) are the subjects of high-throughput studies that make up the bulk of the interactions in curated experimental databases. These viruses happen to have few close relatives (HIV, Influenza A), and infect a limited number of hosts (EBV, HIV), which is why their PPIs are not as readily transferred via orthology as interactions found by text mining for other virus proteins. The viruses that receive the most experimental transfer data are Swine pox virus, Canine oral papillomavirus and Murine cytomegalovirus. The viruses that receive the most text mining transfer data are Gallid herpesvirus, Murine cytomegalovirus and Equine herpesvirus 2.

More than half (55%) of pre-transfer evidence relates to human viruses. However, evidence transferred to human comprises only 26% of all transfered experimental evidence and 18% of all transferred text mining evidence, which implies that the majority of transferred evidence is to a new host (case c or d in figure 1). This is due to the fact that gene duplication events occur less frequently in viruses compared to their host organisms [33], and additionally because fewer species from the virus taxonomic tree have been sequenced and analyzed compared to their hosts [34]. In all, this makes potential transfer partners rarer for transfer between viruses than between hosts.

The distribution of interactions for the 20 viruses with the most interactions is shown in figure 2. The viruses with the largest number of intra-virus interactions include the relatively large double-stranded DNA Herpesvirales and well studied RNA viruses including Influenza and HIV. The same viruses also show the highest proportion of interactions from the experimental channel. An example of two viruses that share interactions based on orthology transfer are human and murine cytomegalovirus (HCMV and MCMV respectively). The majority of the evidence for HCMV is direct evidence, and conversely, the majority of evidence for MCMV is evidence from transfer, which has come from interactions with HCMV.

**Figure 2:**
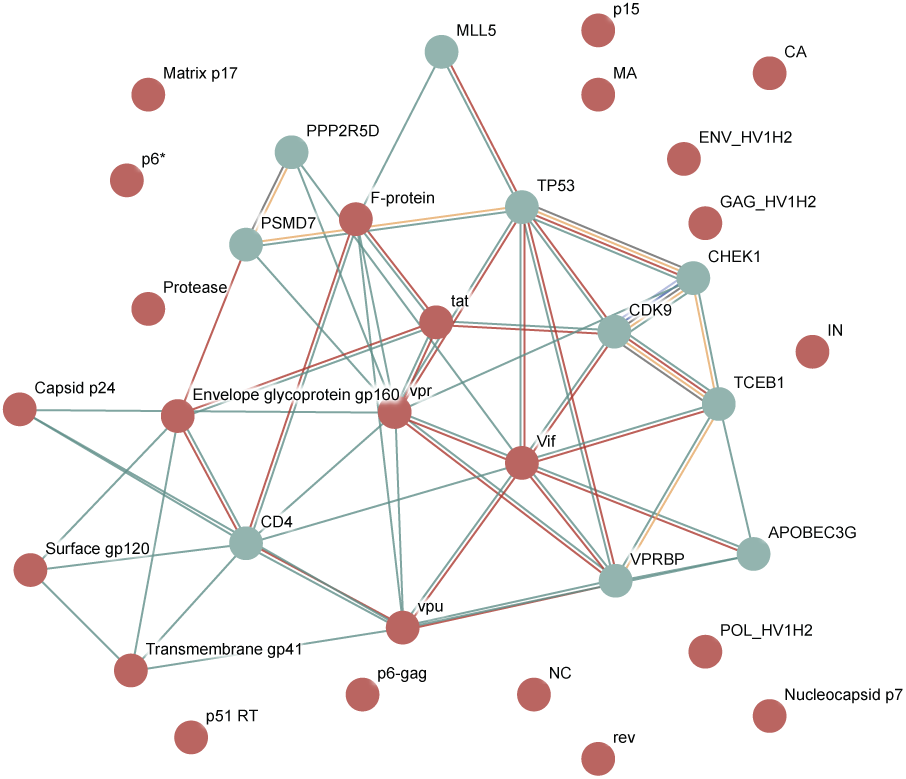
Interactions in viruses.STRING by species (A) Distribution of experimental (red) and text mining (green) interactions, further divided into direct (dark colours) and transferred (light colours) evidence. Data shown for the 20 viruses with the most evidence. Evidence is counted as interaction pairs per channel, such that an interaction that is supported by 3 channels will be counted as 3 evidences. (B) Number of evidences normalized by the number of proteins coded for by that virus.

The virus-virus and virus-host PPI networks are made publicly accessible as a resource which is available at viruses.string-db.org. The data can be browsed online, downloaded from the website, or accessed through the REST API. Further, the data can also be imported into Cytoscape directly [35] using the STRING Cytoscape app [22].

## Utility and Discussion

### Web interface

The Viruses.STRING website enables three variants of protein search: for the complete set of proteins in a virus, for a single protein in a virus, or for multiple proteins in a virus. Since most viral genomes encode only a small number of proteins (the viruses included in the database have a median of 9 proteins), they can easily be displayed in a network together with the most strongly interacting host proteins.

The network interface has a similar appearance to STRING, but the visual styling has been modified to be more flat. The nodes in the network are coloured only based on their origin, either as viral proteins (brick red) or as host proteins (blue-green slate).

As is possible on the main STRING site, the viruses.STRING web interface provides more information about each protein, which is accessed by clicking on the node. Similarly, clicking on any edge displays a summary of the information that contributes to that interaction, and provides links to further inspect the evidence from each channel. Text mining evidence shows highlighted phrases from relevant publications, whereas experiments evidence shows the specific database and publication from which it was obtained.

### Example: HIV-1

In this example, we will query for all proteins present in Human Immunodeficiency virus type 1. If the host field on the search page is left empty, the server will auto detect the host species with the most interactions with the specified virus, in this case, human. An interaction network will then be shown for the virus proteins and for the 10 human proteins that have the highest interaction scores with these virus proteins, as in figure 3. Interaction scores have a cut-off of 0.4 by default, the same as the main STRING site.

**Figure 3:**
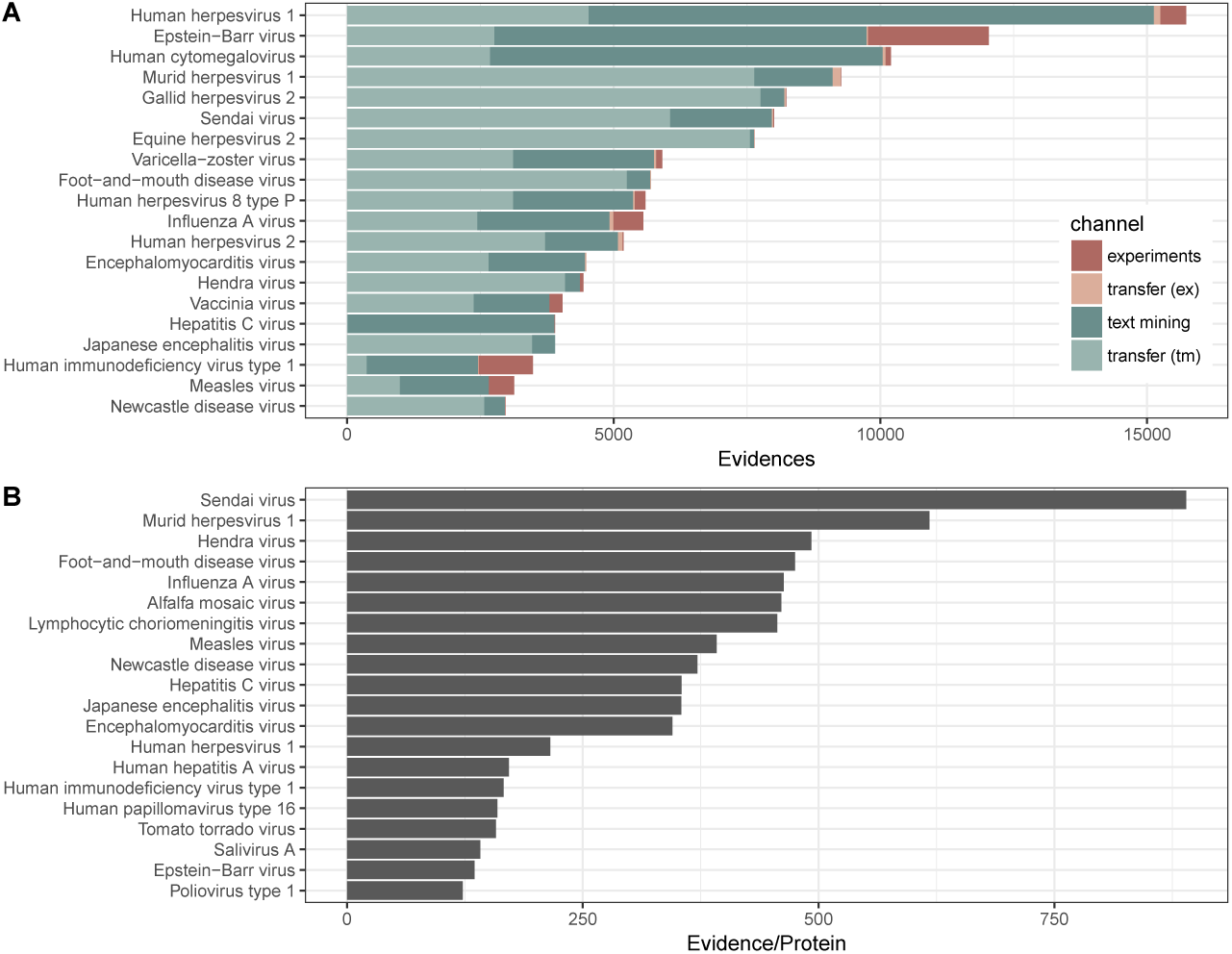
HIV-1 and Homo sapiens interaction network in viruses.STRING HIV-1 and Homo sapiens interaction network downloaded as a vector image from viruses.STRING.

HIV-1 consists of 19 proteins, 10 of which are cleaved from 3 polyproteins. The polyproteins are translated as a single long protein, and then the long polyprotein is cleaved by the viral protease into functional protein units. The database includes 24 proteins as it includes some partial cleavage products, such as both gp160 and gp120 which is cleaved from gp160.

### Cytoscape STRING app

The Viruses.STRING interaction data can also be queried from the Cytoscape STRING app. This requires version 3.6 of Cytoscape or greater and version 1.4 of the STRING app or greater, which is available for free in the Cytoscape app store (http://apps.cytoscape.org/apps/stringapp).

The STRING app allows for more flexible queries than the Viruses.STRING website, such as choosing specific additional host proteins to be included in the network, and displaying multiple hosts and multiple viruses in the same network. In addition to the Viruses.STRING interaction data, the app automatically fetches node and edge information, which can be used for further analysis. The former includes the protein sequence for host and virus nodes, subcellular localization data from the COMPARTMENTS database for human proteins, and tissue expression data taken from the TISSUES database for human, mouse, rat and pig proteins [36]. Edge information includes the combined confidence score from all channels as a probability that the interaction is true.

Figure 4 illustrates the combined interaction network for HPV 16 and HPV 1a proteins with the top 50 human proteins they interact with ranked by combined interaction score. Since HPV is known to disrupt the cell cycle [37], many of the proteins that interact with E6 and E7 are associated with the nucleus and the GO term for cell cycle. A tutorial to reproduce this network in Cytoscape is available at http://jensenlab.org/training/stringapp/.

**Figure 4:**
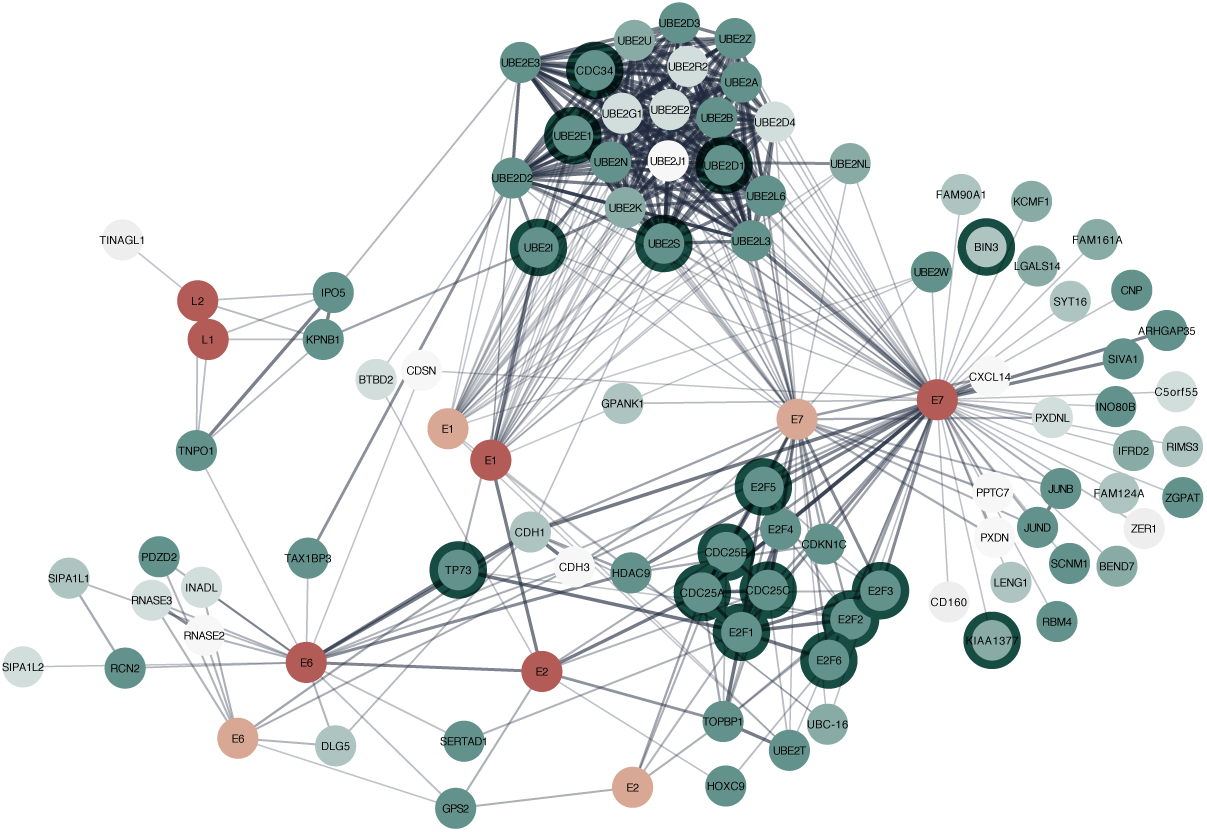
PV and Homo sapiens interaction network in Cytoscape Proteins from Human Papillomavirus type 16, and HPV type 1a with their human protein interaction partners. Virus proteins are coloured according to their species (dark red: HPV 16, light red: HPV 1a). The human proteins are coloured in shades of green with darker colours showing a stronger association with the nucleus. The dark halos around human proteins are those that are associated with the GO term for cell cycle. The HPV E6 and E7 proteins are known to interfere with the cell cycle. This analysis shows some of the data exploration and visualization flexibility that is easily possible within Cytoscape.

## Conclusions

The Viruses.STRING database provides a single unified interface to virus–virus and host–virus PPIs from text mining and many experimental sources. With a simple web interface, the database can easily be queried to immediately retrieve the interaction partners for a protein of interest, and the corresponding evidence can be inspected. The Cytoscape STRINGapp, although it requires software to be installed, provides more versatility than the website, and can handle much larger networks — up to at least as large as the human interaction network. This provides the researcher with more opportunities to answer interesting biological questions about viruses and their hosts. For example, the virus–host network could potentially be used to select candidate host proteins as drug targets to inhibit virus infection, possibly by repurposing existing drugs. This approach would likely generate less viral resistance to the drug since the host protein is being targeted, instead of a viral protein that can mutate easily [38, 14].

As this is the first iteration of Viruses.STRING, there are currently some limitations to the data. The virus data is provided only at the species level, with the exception of Dengue types 1–4, even though there is some evidence that different influenza strains show differential protein interactions [39]. This fine grained resolution will be added in a future version for those viruses where sufficient data is available, such as Influenza A.

Text mining reveals many more virus-host PPIs in the literature than have been collected into databases. The text mining gives good precision and recall for virus species, and good precision for virus proteins [28]. However, the method performs less well for virus proteins in terms of recall, meaning that many interactions may still be missed by this approach [28].

Just as a having a broader view of PPIs has provided a deeper understanding of cellular function [40], having a similar understanding between pathogens and their hosts will provide new information to combat clinically and economically relevant viral infections and diseases.

## Declarations

Availability of data and material: The datasets generated and/or analysed during the current study are available for download at viruses.string-db.org/download

Funding: This work was supported by the Novo Nordisk Foundation (grant NNF14CC0001), and by SIB Swiss Bioinformatics Institute and the University of Zurich. The funding agencies had no role in the design, analysis, interpretation of the data or writing of the manuscript.

## Author’s contributions

HC gathered and analyzed the data. ND integrated viruses into the STRINGapp. DS, CvM, LJJ contributed to the design of the study and revised the manuscript. All authors read and approved the final manuscript.

## Acknowledgements

The authors would like to thank John ‘Scooter’ Morris for his continued work on the Cytoscape STRING app to support these changes. HC would like to thank the members of the Von Mering group for their hospitality during the summer over which the bulk of this work was conducted. This work was supported by the Novo Nordisk Foundation (grant NNF14CC0001) (HC, ND, LJJ), SIB Swiss Bioinformatics Institute and the University of Zurich (DS, CvM).

